# The impact of 1/*f* activity and baseline correction on the results and interpretation of time-frequency analyses of EEG/MEG data: A cautionary tale

**DOI:** 10.1101/2020.12.04.412031

**Authors:** Máté Gyurkovics, Grace M. Clements, Kathy A. Low, Monica Fabiani, Gabriele Gratton

## Abstract

Typically, time-frequency analysis (TFA) of electrophysiological data is aimed at isolating narrowband signals (oscillatory activity) from broadband non-oscillatory (1/*f*) activity, so that changes in oscillatory activity resulting from experimental manipulations can be assessed. A widely used method to do this is to convert the data to the decibel (dB) scale through baseline division and log transformation. This procedure assumes that, for each frequency, sources of power (i.e., oscillations and 1/*f* activity) scale by the same factor relative to the baseline (multiplicative model). This assumption may be incorrect when signal and noise are independent contributors to the power spectrum (additive model). Using resting-state EEG data from 80 participants, we found that the level of 1/*f* activity and alpha power are not positively correlated within participants, in line with the additive but not the multiplicative model. Then, to assess the effects of dB conversion on data that violate the multiplicativity assumption, we simulated a mixed design study with one between-subject (noise level, i.e., level of 1/*f* activity) and one within-subject (signal amplitude, i.e., amplitude of oscillatory activity added onto the background 1/*f* activity) factor. The effect size of the noise level × signal amplitude interaction was examined as a function of noise difference between groups, following dB conversion. Findings revealed that dB conversion led to the over- or under-estimation of the true interaction effect when groups differing in 1/*f* levels were compared, and it also led to the emergence of illusory interactions when none were present. This is because signal amplitude was systematically underestimated in the noisier compared to the less noisy group. Hence, we recommend testing whether the level of 1/*f* activity differs across groups or conditions and using multiple baseline correction strategies to validate results if it does. Such a situation may be particularly common in aging, developmental, or clinical studies.

## Introduction

The study of rhythmic patterns of neural activity, or neural oscillations, has been a central area of interest in neuroscience since their discovery at the dawn of electroencephalography (EEG) research (see e.g., Buzsáki & Draguhn, 2004; Stone & Hughes, 2013). Neural oscillations of various frequencies have been linked to virtually all aspects of cognition, from perception to memory and attention (Başar et al., 2000; Buzsáki, 2002; Buzsáki & Draguhn, 2004; Cavanagh & Frank, 2014; Clayton et al., 2015; Cohen, 2014a; Gratton, 2018; Hanslmayr et al., 2011; Lundqvist et al., 2016; Mathewson et al., 2011, 2009). Methods to identify such narrowband oscillations typically involve transforming neural data to the frequency or time-frequency domain, and locating peaks in the frequency spectrum or increases in power limited to an area of the time-frequency map, respectively (Cohen, 2014b; Herrmann et al., 2014). Importantly, neural data in these domains exhibit a 1/*f*-like power spectrum, that is, amplitude – and thus, power – decreases as a function of frequency following a power-law function (He, 2014). This prominent feature of the frequency spectrum most likely reflects irregular, asynchronous firing of neurons or neuronal assemblies with no periodicity, clearly distinguishing it from rhythmic, oscillatory neural activity (He, 2014; He et al., 2010; Miller et al., 2009, 2014).

While broadband activity resulting from non-oscillatory sources is often referred to as 1/*f* “noise,” it is likely to have functional significance for behavioral performance (Clements et al., 2020; He et al., 2010; Miller et al., 2014; Ouyang et al., 2020). Nevertheless, we will use the terms “1/*f* activity” and “1/*f* noise” interchangeably throughout this manuscript to refer to broadband activity, simply because in studies of narrowband oscillatory activity, this feature is not typically of interest. In fact, in these studies, 1/*f* noise needs to be corrected for in the frequency domain (Demanuele et al., 2007; Donoghue et al., in press; Ouyang et al., 2020; Wen & Liu, 2016) and the time-frequency domain (Cohen, 2014b; Herrmann et al., 2014) because it could otherwise confound estimates of narrowband power, obscure differences in oscillatory activity between frequency bands, and/or create illusory differences where none exist.

As noted above, one way of identifying narrowband phenomena in the brain is to convert electrophysiological data to the time-frequency domain, a method that has gained widespread popularity in the past two decades (Cohen, 2014b). This type of analysis allows us to examine changes in the frequency composition of the data as a function of time, e.g., before or after a stimulus is presented. In studies using this approach, one of the most common methods used to correct for the pervasive 1/*f* power scaling is to baseline correct raw power values (expressed in μV^2^) by dividing each time point by the mean activity in a (typically pre-stimulus) baseline period at each frequency. The resulting values are then converted to decibels (dB) by taking their logarithm (with base 10) and multiplying it by 10 (Cohen, 2014b; Herrmann et al., 2014). These steps are summarized by the following formula:

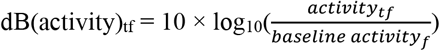

where *t* denotes the time of the activity, and *f* denotes a given frequency. In other words, when dB conversion is applied to time-frequency decomposed neural data, the data undergo two non-linear transformations: first they are divided by a baseline spectrum, then they are converted to a logarithmic scale, operations that are identical to first transforming power to the logarithmic scale and then subtracting the baseline values from the resulting matrix. The log-transformation is applied because power values are strongly positively skewed, and a logarithmic transform of the power values yields distributions that are more symmetrical and therefore more closely approximate a normal distribution. These, in turn, are desirable characteristics for testing hypotheses about changes in the expected value of the power as a function of the variables of interest.

In the present paper we will argue that the first of these two steps (i.e., of division by baseline and log transform) rests on problematic assumptions and that both steps may lead to the distortion of the narrowband signal that researchers typically aim to isolate, following correction for 1/*f* scaling (Demanuele et al., 2007; Donoghue et al., in press; Ouyang et al., 2020; Wen & Liu, 2016), in large part because they involve non-linear operations.

First, using a divisive baseline assumes that the relationship between the power of narrowband oscillations (e.g., a burst of theta oscillations post-stimulus) and broadband background activity with a 1/*f* slope (ubiquitous neural noise) is *multiplicative* rather than additive. In other words, this procedure implies that narrowband oscillatory activity increases or decreases in power proportionally to the baseline activity in that frequency range. This may be a reasonable model when baseline activity is only reflecting a different level of the same narrowband activity, and the experimental manipulation is expected to change it in a multiplicative fashion (i.e., by doubling or halving this baseline level of activity; Delorme & Makeig, 2004). However, this assumption is inappropriate if the baseline and the experimentally-critical levels of activity at any particular frequency are determined by two *separate* phenomena (i.e., if narrowband oscillations and 1/*f* noise are largely uncorrelated). In other words, unless these two phenomena reflect different levels of activation in the same underlying source, even in the presence of experimental manipulations, a division by the baseline value (i.e., the multiplicative model) will provide inaccurate results.

Furthermore, if we accept the idea that non-oscillatory (1/*f*) activity is contributing to the spectrum (He, 2014), it is entirely possible that for some frequencies such activity could account for practically all the observed power during the baseline period. In such cases, it becomes difficult to conceptualize that increases in narrowband activity at that same frequency represent a proportional amplification of ongoing narrowband oscillations, as no such oscillations may be present at all, or be so small as to be very difficult to measure.

A model where variance associated with independent factors is additive can be a viable alternative to the multiplicative model. This additive model assumes that there is a high degree of independence between the changes induced by experimental manipulations on narrowband and broadband (1/*f*) activity. As the variance of a signal corresponds to its power, the power of these factors would therefore also be additive. In terms of neural signal generation, such an additive model would assume that oscillatory activity is added on top of pre-existing background activity and is thus largely independent of the level of such background activity (Grandchamp & Delorme, 2011).

Using empirical data, in the first part of this paper we will investigate whether the level of broadband activity and the magnitude of narrowband activity in the alpha range (8-12 Hz) scale together during rest, and are therefore strongly positively correlated as predicted by a multiplicative model or are instead independent, and therefore largely uncorrelated, in line with an additive relationship. Foreshadowing our results, we find no robust positive relationship within participants between broadband activity (identified here with the scaling factor of the 1/*f* noise) and power in the alpha range after detrending the data by subtracting the effects of 1/*f* activity in the alpha band. As such, dividing by baseline power in time-frequency analyses may be ill-advised, because the same signal could appear to be different following baseline correction depending on the level of noise present at that given frequency due to independently varying 1/*f* activity. For instance, a narrowband signal of magnitude 2 μV^2^ in the presence of 5 μV^2^ broadband noise would have a corrected effect size of (2+5)/5 = 1.4, whereas with 8 μV^2^ broadband noise the corrected effect size would be (2+8)/8 = 1.25, and so on.

Importantly, this noise-dependent distortion may be further exacerbated by the next step of dB conversion, logarithmic transformation. The logarithmic function is steeper for lower as opposed to higher values, meaning that low values are amplified while high values are dampened. There are mathematical consequences to this log-transformation property, which will occur when narrowband (e.g., alpha or theta activity) and broadband (e.g., 1/*f* noise) activities are largely uncorrelated. Specifically, for the same amount of change from baseline in narrowband activity following baseline division, the power change values for a specified frequency band will be numerically smaller if the background broadband activity noise is high compared to when it is low. In other words, the same difference between two baseline corrected values - (signal + noise)/noise-will appear to be gradually smaller as the denominator of this ratio (i.e., 1/*f* noise) increases. This is illustrated in **Figure 1**. The X-axis shows 50 proportion values where the signal was fixed to be 2 μV^2^ and the added noise ranged from 1 to 20 μV^2^. The log-transforms of the values following baseline division are plotted on the Y-axis. As expected, log-transformation clearly accentuates differences between lower values and attenuates differences between higher values in a non-linear fashion. Taking all this into consideration, dB conversion could create problems in situations where the same change in a signal is compared between conditions or groups with differing levels of noise: higher noise will lead to smaller values following division compared to lower levels of noise, which will move these values closer to the steeper end of the curve as opposed to the flatter part, accentuating differences.

**Figure 1.**
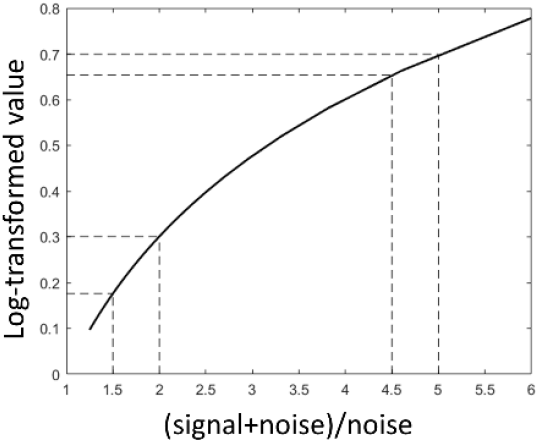
A series of (noise ± signal)/noise ratios and their corresponding log10 values. Dashed lines illustrate how log-transformation changes the difference between values as a function of the values themselves. Signal was always 2 arbitrary units (AU), while noise increased from 1 to 20.

In the main part of the present paper we demonstrate how distortions in time-frequency data caused by dB conversion complicate the interpretation of findings when noise and signal are additive. To this end, we ran a series of simulations, modelling a mixed-design with a within-subject factor (Condition 1 vs. Condition 2) and a between-subject factor (Group 1 vs. Group 2). This type of design is common in individual difference research, including aging, developmental, and clinical studies. A time-limited narrow-band oscillatory signal was added onto broadband random noise that displayed 1/*f* scaling in the frequency domain, in each condition and for each group. To explore the impact of different amounts of 1/*f* noise in different groups on the results of time-frequency analysis, we increased the level of broadband activity in one group by increasing the scaling factor of the 1/*f* function^1^, across iterations of the simulation, while keeping the 1/*f* scaled broadband activity constant in the other group. As such, the level of noise was fixed in one group (fixed-noise group), and variable in another (variable-noise group), changing from iteration to iteration. Three different scenarios were simulated: 1) the difference between Condition 1 and Condition 2 is the same in both fixed-noise and variable-noise groups (no Group × Condition interaction); 2) the difference between Condition 1 and Condition 2 is greater in the fixed-noise group than in the variable-noise group (Group × Condition interaction, fixed > variable), and 3) the difference between Condition 1 and Condition 2 is smaller in the fixed-noise group, than in the variable-noise group (Group × Condition interaction, fixed < variable). The main objective was to determine whether the size of the interaction effect changed as a function of the noise difference between the two groups, i.e., as the level of broadband activity increased in the variable-noise group compared to the fixed-noise group, when dB conversion, simple baseline subtraction, or no correction were applied to the data. We expected to see a greater noise-dependence of the effect size following dB conversion for the reasons outlined above. Specifically, the effect size in the variable-noise group is likely to be gradually underestimated as a function of broadband activity (i.e., as noise increases) potentially leading to the appearance of fixed > variable interactions even when none are present.

Notably, this design was chosen because the 1/*f* property of the power spectra of neural data has recently been shown to change with age (Clements et al., 2020; Dave et al., 2018; Voytek et al., 2015), clinical diagnosis (Robertson et al., 2019), and behavioral states (Miller et al., 2014; Podvalny et al., 2015). These findings make the modelled situation, whereby a within-subject change (e.g., the difference between one condition that purportedly engages a cognitive process and one that does not, such as the incongruent and the congruent conditions in a conflict task) is compared between two groups that differ in broadband activity (e.g., younger and older adults) highly plausible and in fact, common in typical analyses of group differences. As such these findings are relevant to most electrophysiological (and electromagnetic) studies of individual and group differences, including those common in aging, developmental and psychopathological research.

## Methods

### Participants

To investigate our first research question, related to the additivity of narrowband and broadband signals (i.e., whether fluctuations in broadband activity are correlated with fluctuations in narrowband activity across time) we used resting state data from two previous lab projects, comprising 80 participants (mean age = 40 ± 24, 56.25% females), and two age groups (N_1_ = 51, 18-31 year-olds and N_2_ = 29, 65-83 year-olds). No participant reported any major health issues, including psychiatric or neurological conditions, and all had normal or corrected-to-normal vision and hearing.

To investigate our second research question (i.e., how dB conversion distorts time-frequency data when signal and noise are additive) we ran a series of simulations, described below.

### Procedures

#### Broadband-narrowband correlations

EEG and EOG were recorded from 64 channels using BrainAmp amplifiers (BrainVision Products GmbH) while participants were instructed to sit without moving, with their eyes open for a little over 1 minute, then with their eyes closed for a little over 1 minute.

EEG was recorded referenced to the left mastoid, and subsequently re-referenced to the average of the two mastoids off-line. Sampling rate was 500 Hz, and impedance was kept below 10kΩ. During pre-processing, data were segmented into epochs of 4096 ms. Ocular artifacts were corrected using the method described in Gratton et al. (1983). Epochs with voltage fluctuations exceeding 200μV were excluded from analyses. Data were subsequently low-pass filtered at 30 Hz using a Hamming windowed *sinc* finite impulse response filter as implemented in the pop_eegfiltnew() function of EEGLAB (Delorme & Makeig, 2004).

The remaining epochs were then subdivided into shorter ∼1000 ms segments to increase the number of observations per participant. Then, for each segment in each condition (eyes open or eyes closed), the offset of the 1/*f* slope and power in the alpha frequency range (8-12 Hz) were identified at electrode Pz. To do this, first the fast Fourier transform of the time-domain data was calculated using the fft() function in MATLAB 2019b (The MathWorks, Inc., Natick, Ma, USA). Frequencies below 2 Hz and above 25 Hz were discarded from further analysis. For the 1/*f* calculations, frequencies between 4 and 20 Hz were also removed so the estimation of broadband characteristics would not be contaminated by narrowband oscillations potentially present in the theta to beta range. A linear model of the form *power* = ß_0_ + ß_1_(1/*f*) was fitted to the remaining frequencies. In this formalization of 1/*f* scaling, the exponent of the slope is assumed to be -1, and the ß_1_ coefficient reflects the offset (scaling factor) of 1/*f* activity. In log-log space, this parameter is roughly equivalent to the intercept of the regression line and can be interpreted as the level of broadband activity on a given trial. As such, the offset parameter of each segment was saved for further analyses. Using the ß_0_ and ß_1_ coefficients generated by the model, a regression line was fit to the full spectrum from 2 to 25 Hz and subtracted from the data to remove 1/*f* activity. From this detrended spectrum, the mean power between 8 and 12 Hz was extracted, this constituted our measure of alpha power. Once this has been done for all segments in a condition, Kendall’s τ was calculated between the 1/*f* beta weights (offset values) and the mean alpha values across segments within a subject to examine the relationship between the level of broadband activity and the magnitude of narrowband signals. These τ values were then saved for each participant.

We chose to analyze resting state EEG data to ensure that any relationship that emerges between broadband and narrowband activity is not due to task-induced changes that affect both components of the spectrum separately, possibly creating illusory correlations, which could be either positive or negative. Please note that even during rest it cannot be entirely ruled out that any correlation that emerges between the two variables is due to some underlying change that affects both simultaneously (e.g., a shift in mental state); however, since no systematic task was given, the possibility of such a spurious correlation was minimized.

#### The effects of dB conversion

Simulations were run in MATLAB 2019b to investigate the effects of dB conversion on time-frequency data at various levels of noise. Two groups were simulated (Groups 1 and 2) both with 30 participants, and with 200 trials (epochs) for each participant, 100 each for Condition 1 and Condition 2. The difference between the two groups and the two conditions is detailed below. The generated epochs were 4096-ms long and the sampling rate was set to 500 Hz. For each epoch, first white noise was generated in the time domain, which was then converted to the frequency domain using the fft() function in MATLAB. The resulting amplitude values were then multiplied by the slope of a curve that was originally fit to the logarithm of the frequencies making up the spectrum. This curve had an intercept of 0 and an exponent of −0.5. This exponent value was chosen to make sure that the slope of the 1/*f* function would be approximately −1 when power (amplitude^2^) values are considered as opposed to amplitude values, as is typical in the literature. The resulting weighted amplitude spectrum was then transformed back to the time-domain via the inverse Fourier transformation as implemented by the ifft() function in MATLAB. The level of noise, i.e., the offset of 1/*f* activity, was modulated by multiplying this time-domain signal by a value between 3 and 15 with 50 linearly spaced values covering this range. In the fixed-noise group, this number was always the lowest number, 3. In the variable-noise group, the number increased on each iteration of the simulation. Each of these 50 iterations was run 10 times, and results were averaged across runs, separately for each noise level.

The narrowband signal was simulated by adding a wavelet to the noise in the time-domain. This wavelet constituted the signal of interest and was created by multiplying a 5-Hz sine wave with a Gaussian taper equal to the length of the epoch. The center of the wavelet was at 2148 ms. Random Gaussian noise was added to the timing, duration, and phase of the signal on each trial to simulate random intraindividual variability seen in real recordings. Three scenarios were modelled (see **Table 1**), so that the 10×50 iterations were run three different times. In scenario one, the amplitude of the signal was 1 in Condition 1 and 1.2 in Condition 2 for both the fixed-noise and the variable-noise group. In other words, the difference between conditions was the same in both groups, i.e., there was no Condition × Group interaction. Consequently, we will refer to this scenario as the no Group × Condition interaction. In scenario two, amplitudes were 1 and 1.2 for Conditions 1 and 2 respectively in the variable-noise group, but 1 and 1.3 in the fixed-noise group (Group × Condition interaction, fixed > variable). Finally, in scenario three, the signal difference was bigger in the variable noise group (1 vs. 1.3) than in the fixed-noise group (1 vs 1.2; Group × Condition interaction, fixed < variable).

**Table 1.** Parameter values used in the simulations investigating the effect of dB conversion on time-frequency decomposed data.

| Parameter | Scenario 1 |  | Scenario 2 |  | Scenario 3 |  |
| --- | --- | --- | --- | --- | --- | --- |
|  | Fixed-noise | Variable-noise | Fixed-noise | Variable-noise | Fixed-noise | Variable-noise |
| Noise level | 3 | 3-15 | 3 | 3-15 | 3 | 3-15 |
| Condition 1 | 0.10 | 0.10 | 0.10 | 0.10 | 0.10 | 0.10 |
| Condition 2 | 0.14 | 0.14 | 0.17 | 0.14 | 0.14 | 0.17 |

Note: The values for Conditions 1 and 2 represent the maximum expected power values following the time-frequency decomposition of the signal. These maximum values were calculated by running time-frequency analyses on the wavelet signals without any added noise (including 1/f activity). Fixed-noise = fixed-noise group, variable-noise = variable noise group. For the description of Scenarios 1-3 see text.

In all three scenarios, on each iteration of the simulation (i.e., at each noise level), a time-frequency analysis was conducted on each simulated subject’s data, separately for the two conditions, following methods described in Cohen (2014b). To do this, epoched data were fast Fourier transformed and multiplied by the fast Fourier transform of Morlet wavelets of different frequencies. Morlet wavelets are complex sine waves tapered by a Gaussian, similar to the signal described above. Thirty frequencies were used to create these wavelets, logarithmically spaced between 3 Hz and 30 Hz. The number of wavelet cycles (i.e., the width of the Gaussian taper) increased with frequency from 3 to 10 to adjust the balance between temporal and frequency precision across different frequency bands. Next, the inverse Fourier transform of the product spectra were computed, and power values were then obtained from the resulting complex signal by squaring the length of the complex vector at each time point (Cohen, 2014b). Power values were averaged across trials, then the first and last 500 ms of the epochs were removed to avoid edge artefacts.

The resulting time-frequency matrix was then processed in two alternative ways: 1) mean activity in the baseline period was subtracted from all time points at each frequency (subtraction method); and 2) time-frequency points were converted to dBs by dividing each time point at each frequency by the mean activity in the baseline, and then taking 10×log10 of the resulting ratios (dB conversion). The baseline period was defined as the first 500 ms of the epochs after trimming to avoid edge artefacts. The uncorrected, raw power matrices were also used in further analyses. (Please note that baseline correction was done for each condition separately, however, this is unlikely to introduce any between-condition differences as the baseline period was not modulated systematically between conditions.)

Following this processing, maximum power was identified within a window stretching ± 20 ms around the time point set as the center of the wavelet signal, and ± 2 Hz around the central frequency used to create the signal for the uncorrected and the two corrected matrices in each condition. Then, power was averaged within a smaller window stretching ± 10 ms and ± 1 Hz around the time point and the frequency of the peak value, respectively, to create the power values used in statistical analyses for each subject^2^. We will refer to these analyses as “pseudo-confirmatory analyses” as they model a situation in which the researcher has a clear hypothesis about the timing and the frequency range of the effect. For each iteration of the simulation, the difference between the mean values for Condition 1 and Condition 2 was calculated for each participant in each group. Then the effect of Group (fixed-vs. variable-noise) on this difference variable was tested using an independent sample *t*-test. Mean differences and *t*-values are reported as a function of noise difference between groups and correction method.

To model a situation where the hypothetical researcher is more naïve regarding the characteristics of the effect of interest, “pseudo-exploratory analyses” were also run using permutation testing. On every iteration of the simulation, the time-frequency matrix of Condition 1 was subtracted from the time-frequency matrix of Condition 2 for each individual in both groups. Then 1000 permutations were run, randomly shuffling the Group label on each difference matrix, and subsequently calculating the difference between the mean difference matrix of “Group 1” and the mean difference matrix of “Group 2”. The mean and standard deviation of the resulting 1000 difference-of-differences matrices was then used to standardize the real Group 1 – Group 2 difference map based on the observed data. Any points with an absolute *Z*-value smaller than 1.6449 - corresponding to an α of .05 – were set to 0. A cluster-based correction for multiple comparisons was then used to identify significant effects. Each permutated matrix was similarly thresholded, and a null-distribution of cluster sizes was created by identifying the number of pixels in the largest cluster – i.e., contiguous pixels with above-threshold *Z*-values – in each matrix. Any clusters in the original thresholded map based on the observed data that were larger than the 95^th^ percentile of the null distribution were labeled significant; all other clusters were disregarded. The mean *Z*-values within these clusters will be reported for each baseline correction method and at each level of noise.

## Results

Data and code for all analyses can be found at the following URL: https://osf.io/sf8pv/.

### Broadband-narrowband correlations

**Figure 2A** shows the distribution of Kendall’s τ values between 1/*f* offset and detrended alpha power across participants at electrode Pz in the eyes closed condition where alpha power was the highest^3^. The mean number of data segments across participants was 75.65 ± 14.76. The mean of the correlation values was overall slightly negative as opposed to positive as would be predicted by the multiplicative model of signal generation. We reasoned the negative correlation occurs because the estimated 1/*f* slope is likely to vary around the real slope. If observed power is the sum of 1/*f* activity and a narrowband signal in the alpha range, subtracting an over-/underestimate of the former from the observed power will lead to an under-/overestimation of the latter resulting in a negative correlation.

**Figure 2.**
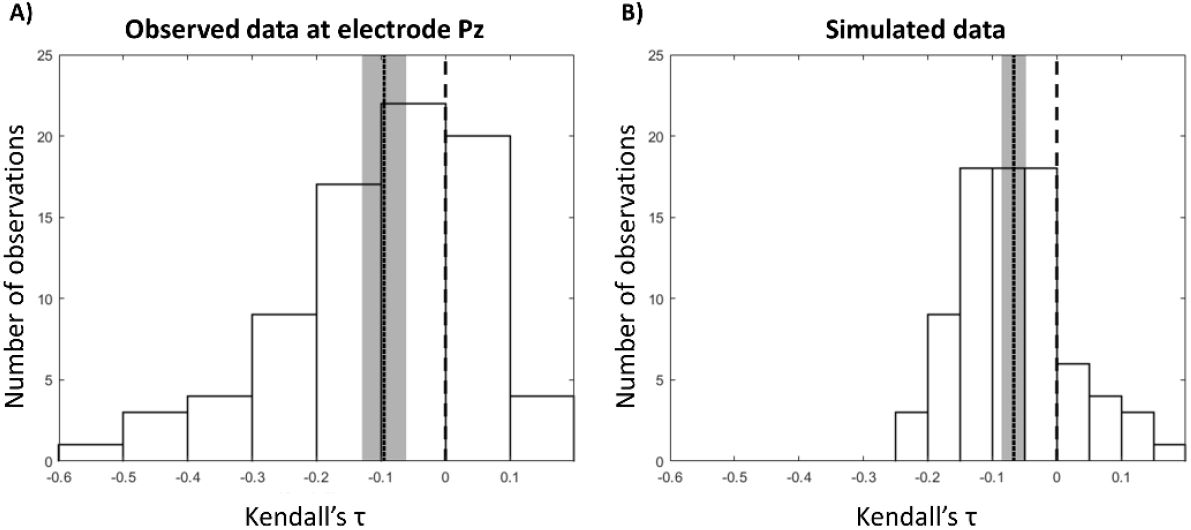
Distribution of Kendall’s τ values across participants, quantifying the relationship between within-subject fluctuations of level of 1/f activity and magnitude of alpha power in real resting state data (A), and simulated data (B). The dotted line is the mean correlation across participants, the shaded area is the 95% confidence interval around the mean. Dashed line indicates 0.

To test this idea, we ran simulations with 80 participants, and 75 observations per participant. Each simulated epoch was the summation of 1/*f*-scaled noise and a 12 Hz wavelet (both generated the same way as described in the Procedure section for the main simulations). On each 1024-ms long trial, the magnitudes of noise and signal were modulated independently by multiplying each in the time domain by a random number. The same method was used on these data to determine within-subject broadband-narrowband correlations as on the real data. When noise and signal were modulated independently, a small negative correlation emerged just like in the real data (**Fig. 2B**). When noise and signal were yoked, i.e., noise was multiplied by a random number on each trial, and signal was multiplied by the same number divided by a constant, a positive correlation occurred. As such, if noise and signal scaled together in the real data as would be expected based on the multiplicative model, we should have observed a positive correlation. The small negative correlation between narrowband (alpha) and broadband (1/*f*) activity we did observe was therefore likely artefactual, and consistent with predictions made under the hypothesis of a true zero correlation between these two types of activities, but inconsistent with the predictions made under the hypothesis of a positive correlation between these activities.

### The effects of dB conversion

**Figure 3** shows the most important steps occurring at each iteration of the simulation. The example shows data in the fixed-noise group and the variable-noise group (with the highest possible level of broadband activity) from the time domain through the time-frequency domain to the final time-frequency map, showing the between-group difference of the conditional differences, i.e., the Group × Condition interaction map, in the fixed < variable situation (Scenario 3).

**Figure 3.**
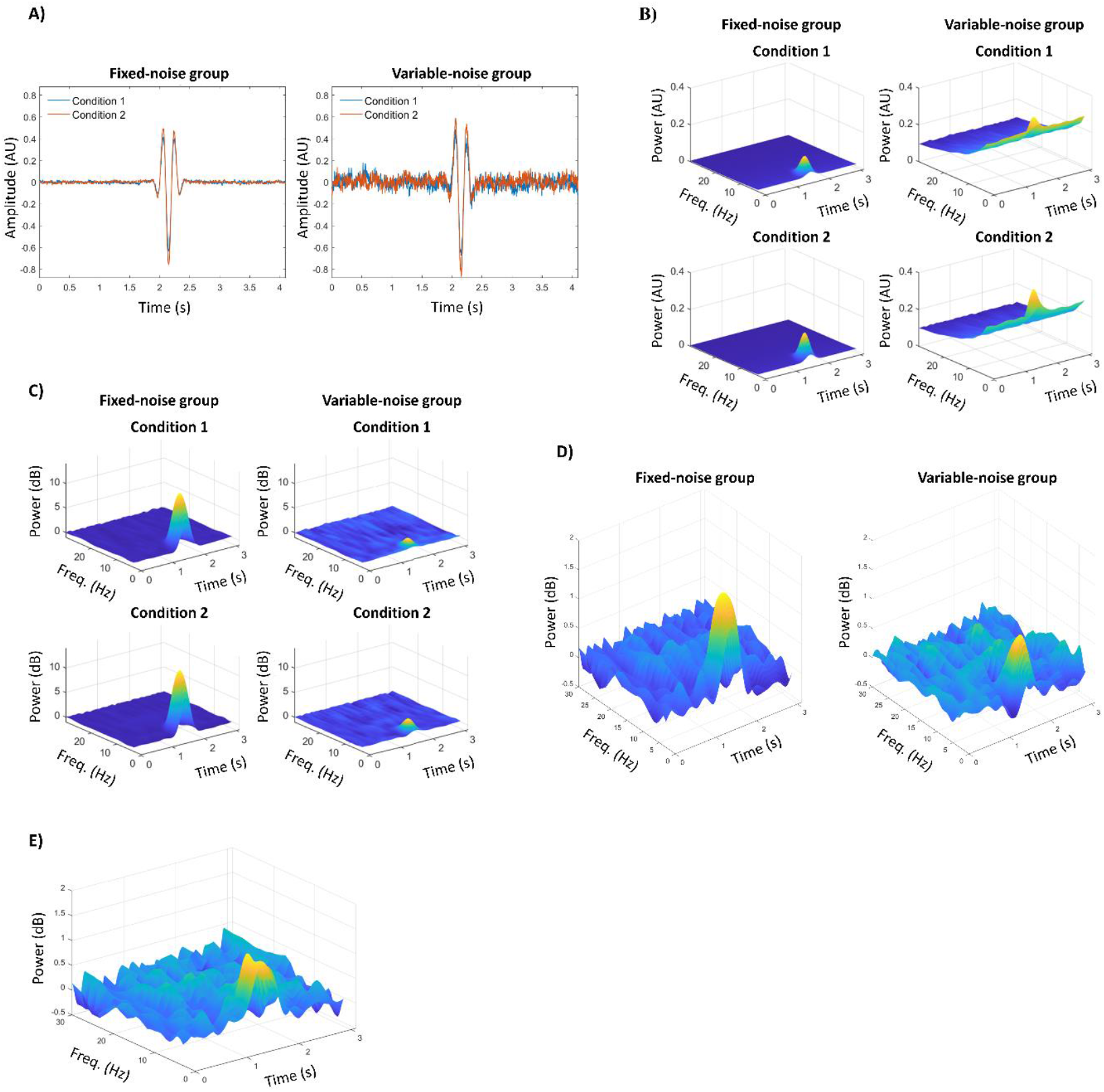
One iteration of the simulation. In this example, the variable-noise group has the highest possible level of background activity (i.e., this would be the final, 50th iteration of a run). The scenario is Group × Condition interaction (fixed < variable). Only dB correction is shown, but uncorrected and baseline corrected data were also analyzed in the time-frequency domain (see text). (A) Trial-averaged data in the time domain across the two groups. (B) 3D plots of uncorrected power in the time-frequency domain in each condition in each group. (C) dB-corrected power in each condition and group. (D) Condition 2 – Condition 1 difference map in the fixed-noise and variable-noise groups, using the dB corrected values from (C). (E) Final interaction map calculated as the difference between the two maps in (D).

**Figures 4-6** illustrate the mean group difference of condition differences (interaction effect), mean *t*-values, and mean *Z*-values within significant clusters, for the three Group × Condition scenarios, respectively (see **Table 1** for details), averaged across the 10 runs of the simulation, as a function of noise level in the variable noise group for each baseline correction method. Panels A, B, and C within each figure show the three scenarios: A) no Group × Condition interaction; B) Group × Condition interaction, fixed > variable, and C) Group × Condition interaction, fixed < variable scenario, respectively.

**Figure 4.**
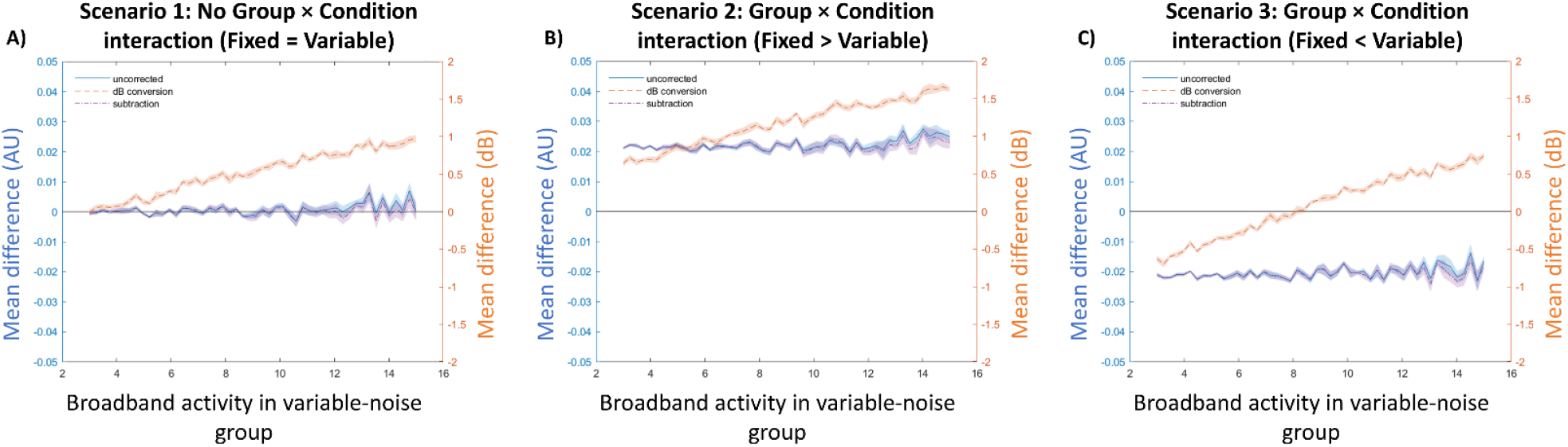
The size of the interaction effect as a function of correction method and noise difference between the two groups. The conditional difference (Condition 2 – Condition 1) in the variable-noise group is subtracted from the fixed-noise group. Uncorrected and baseline-subtracted values are plotted on the left-hand scale, dB corrected values on the right-hand scale. (A) No interaction between group and condition. (B) The conditional difference is bigger in the fixed-noise group than in the variable-noise group. (C) The conditional difference is bigger in the variable-noise group than in the fixed-noise group.

As can be seen, the narrowband activity estimated using the dB correction method is generally more sensitive to the level of broadband activity than the subtraction method. This is because baseline division attenuates signal magnitude as the level of additive noise increases. This noise sensitivity is particularly evident in the no interaction and fixed < variable scenarios (Panels A and C across all figures). In the former scenario, dB conversion led to the emergence of differences that did not exist – both the *t*-tests and permutation testing identified erroneous significant Group × Condition interactions in the fixed > variable direction, even though the within-subject Condition contrast was equivalent for the two groups. This is a consequence of the attenuation of signal magnitude in the variable-noise group as a function of noise. In the fixed < variable scenario, while *t*- and Z-values decreased as a function of noise for all correction methods, they were consistently smaller, and even flipped sign for dB converted values at higher noise levels (**Figs. 5C and 6C**). This suggests that at very high levels of background activity, dB conversion can even reverse the direction of the interaction, as the conditional difference is underestimated in the high noise group compared to the lower noise group. This reversal effect is further illustrated in **Fig. 3** at the most extreme noise difference level, between the fixed-noise group and the variable-noise group. As can be seen, the conditional difference (**Fig. 3D**) is greater in power and larger in area in the fixed-noise (minimal-noise) group compared to the variable-noise (maximal-noise) group following dB conversion. Consequently, in the interaction map showing the results of the fixed-minus-variable subtraction (**Fig. 3E**), there is an increase in power in the frequencies and time points surrounding the central frequency and central time point of the simulated effect, corresponding to an *illusory* fixed > variable interaction effect in the fixed < variable condition. This increase is less pronounced at the exact location of the effect, where the variable-noise conditional difference is largest (**Fig. 3D**), resulting in a truncated cone shape in **Fig. 3E**.

**Figure 5.**
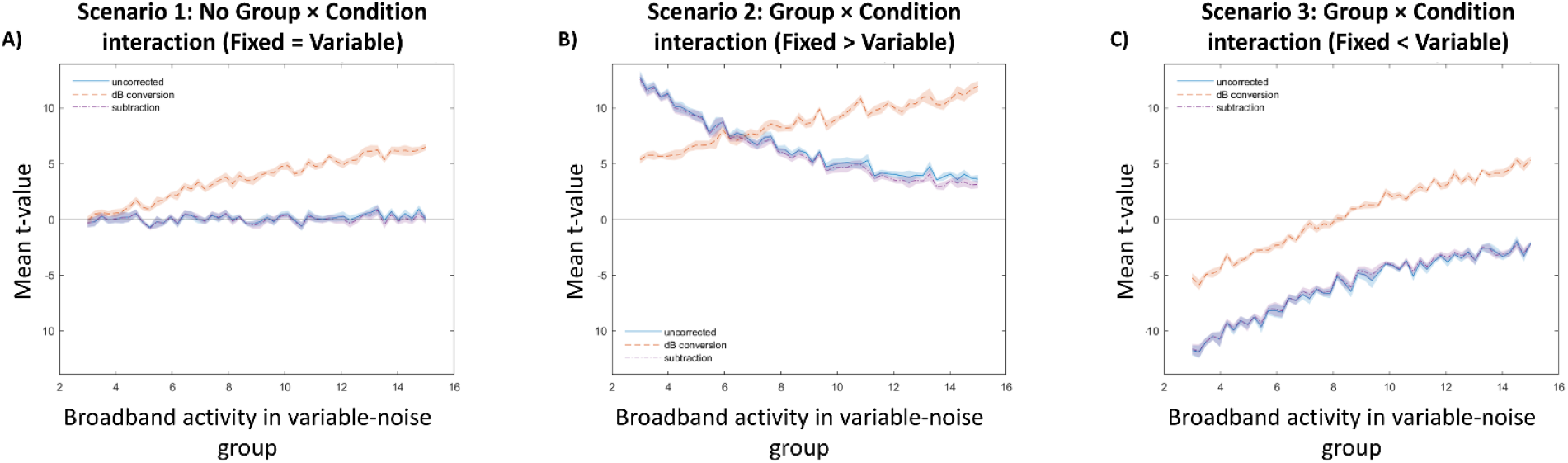
Pseudo-confirmatory analyses. Mean t-values across iterations as a function of correction method and noise difference between the two groups. The conditional difference (Condition 2 – Condition 1) in the variable-noise group is contrasted with the conditional difference in the fixed-noise group using an independent-sample t-test. (A) No interaction between group and condition. (B) The conditional difference is bigger in the fixed-noise group than in the variable-noise group. (C) The conditional difference is bigger in the variable-noise group than in the fixed-noise group.

Finally, findings in the fixed > variable scenario (Panel B in **Figures 4-6**) suggest that dB conversion overestimates the difference between means as a function of noise. This is also reflected in *t*-values that appear to be more stable than those resulting from the other correction methods. The inflated effect size is once again a reflection of the underestimation of the within-subject contrast in the variable-noise group as noise increases.^4^

**Figure 6.**
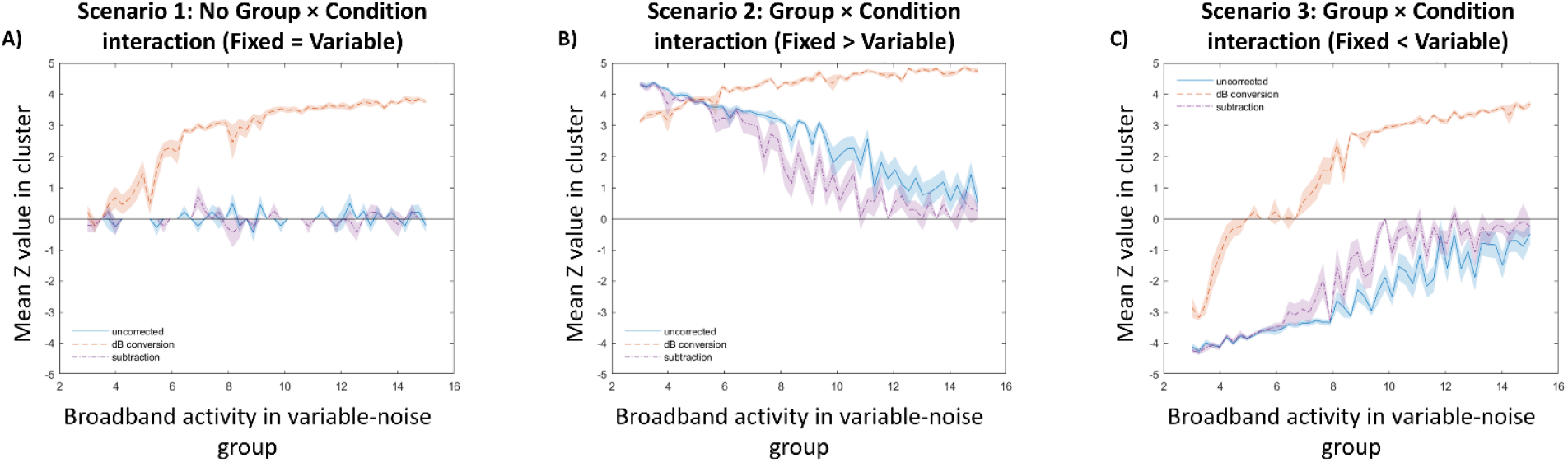
Pseudo-exploratory analyses. Mean Z-values within the significant pixel-based cluster identified by permutation testing in the final interaction map, averaged across iterations, plotted as a function of correction method and noise difference between the two groups. (A) No interaction between group and condition. (B) The conditional difference is bigger in the fixed-noise group than in the variable-noise group. (C) The conditional difference is bigger in the variable-noise group than in the fixed-noise group.

## Discussion

Neural data in the frequency domain contains broadband activity showing prominent 1/*f* scaling, as well as more specific narrowband activity (e.g., alpha oscillations). When the interest is in changes of narrowband activity, power data from time-frequency decomposition of EEG/MEG recordings are often converted to dB values by taking their log10 following baseline division, mostly to reduce the skewing of the power distribution, but also with the intent of removing the effects of broadband phenomena that are often considered to be noise. Implicitly, this method relies on a multiplicative model of signal generation, by which the power ratio of the narrowband and broadband activity for any given frequency is maintained constant across time and conditions.

In the present study, we first offered empirical evidence that a multiplicative model does not adequately describe the relationship between background activity and signal. Estimated 1/*f* offset and alpha power, calculated following the removal of the 1/*f* slope, did not scale together in resting state data, i.e., they were not positively correlated over time within participants. In fact, a small negative relationship was observed; however, simulations indicated that the slight negative relationship might have been the result of subtracting noisy estimates of 1/*f* from the observed power spectra.

Next, using simulated data based on an additive model of signal generation, we illustrated the pitfalls of dB conversion in situations where groups or conditions with different broadband activity are contrasted. Various correction methods were compared, and while they were all sensitive to increases in non-oscillatory activity (i.e., noise) to some extent, this noise-dependence led to unexpected illusory effects for dB conversion, whereas it mostly manifested as a the expected decrease in statistical power for subtraction-based correction.

### Additivity vs. multiplicativity in signal generation

Our finding that 1/*f* offset and alpha power were not positively correlated during rest suggests that a person’s level of broadband activity does not strongly predict the magnitude of their narrowband activity. This is consistent with an additive model of signal generation where narrowband oscillations emerge on a background of random arrhythmic activity, in other words, where signal is added to noise. The multiplicative model, instead, posits that changes in narrowband oscillations are the consequence of the scaling up or down of oscillations already present in those frequency bands as background activity. This would produce a positive correlation between broadband activity and narrowband activity, as higher broadband activity would mean more power across frequencies (especially lower frequencies) which, in turn, would have to result in larger changes in any given frequency when multiplied by a scaling factor.

Interestingly, Muthukumaraswamy & Liley (2018) did observe strong positive within-subject correlations between properties of 1/*f* slope and alpha power. However, these correlations were based on the steepness of the 1/*f* log-log slope (i.e., the exponent of the 1/*f* function, which is a different parameter than that used in the current study, in which the exponent was instead fixed at −1) in the 20 to 100 Hz range, and were not observed when activity at lower frequencies was considered. As such, they are more informative about the relationship between alpha and 1/*f* activity at higher frequencies, which are often ignored in time-frequency analysis, and less so about the relationship between level of baseline activity in the frequency range of the narrowband signals of interest (typically delta to beta ranges – 0.5 to 30 Hz - in human EEG/MEG studies, including the current study). In terms of the physiological interpretation of the different 1/*f-*related parameters, it has been proposed that when local field potentials are measured, the offset of 1/*f* activity represents population spiking, while the steepness of the slope is related to the temporal structure of spiking. However, it is still unclear if these interpretations hold for surface M/EEG (Voytek & Knight, 2015).

Naturally, our findings do not invalidate the claim that some EEG effects may reflect the amplification or dampening of specific narrowband activity – or indeed even of broadband phenomena. For instance, event-related theta-power increases in certain contexts may reflect the amplification of endogenous oscillations that could be occurring even in advance of stimulus presentation (Cohen, 2014a; Cohen & Donner, 2013). In this sense, a hybrid model cannot be ruled out, whereby the frequency spectrum of the baseline, pre-event neural activity is made up of 1/*f*-scaled background activity that is not rhythmic *and* weak narrowband oscillations at different frequencies (additive component). When a stimulus occurs, these weak oscillations are scaled up or down (multiplicative model), or their phases reset to a specific value to generate event-related signals (phase resetting, e.g., Makeig et al., 2002; Sauseng et al., 2007). This hybrid model, however, is not conducive to a simple mathematical analysis approach, such as baseline division and dB transformation, which, as shown here, would introduce substantial distortions to the data. In this sense, an advantage of the additive model is that it avoids making as of yet unsupported assumptions about the relationships between different sources of variance, or power, in the data.

Note that in the current study we have proposed the use of an additive approach, in which the first step for real frequency or time-frequency data may be the estimation of broadband (1/*f*) activity based on frequency bands not of interest for narrowband analyses, followed by subtraction of the broadband effect from all frequencies, and subsequent analysis of the narrowband effect^5^. This stepwise methodology is consistent with work conducted in other labs that also attempt to tease apart broadband and narrowband effects, by subtracting an estimate of the former from the latter (Donoghue et al., in press; Ouyang et al., 2020).

### Decibel conversion in time-frequency analysis

In our simulations, dB conversion led to noise-dependent distortions in the data. Specifically, when an additive relationship between level of background activity and narrowband signal of interest was simulated, conversion to dB scale led to the emergence of illusory interactions between groups, characterized by different broadband activity, and to serious misrepresentations of existing interactions. **Figures 4-6** suggest that we do not have to assume a large difference between two groups in terms of broadband activity for these distortions to appear, i.e., differences between the raw, uncorrected effect sizes, statistical parameters and their dB corrected counterparts were already present at low and intermediate levels of noise, not just at the extreme ends of the scale. This suggests that these anomalies could be present in real data, for instance in aging, developmental, and clinical research when two groups with different levels of neural variability are compared in terms of a within-subject contrast (e.g., pre- and post-intervention performance; low- and high-demand conditions; social observation and no social observation conditions).

Most worrisome, group-by-condition interactions – significant differences in the within-subject contrast between groups – emerged when no actual difference existed between groups in signal magnitude, only in noise magnitude. As with most observed distortions, this is because baseline division reduces the signal in the group with the higher noise level (here the variable-noise group) compared to the lower noise level group (here, the fixed-noise group) resulting in an apparent difference, which then undergoes a non-linear transformation when it is converted to the log scale. Log-transformation accentuates low values, and this could potentially compensate for the reduction in signal magnitude to some extent (**Fig. 1**); however, our findings clearly suggest that this is not sufficient. In practical terms, these distortions could mean that following dB correction of time-frequency data, a researcher could conclude that two groups differ in their oscillatory response to an intervention or in narrowband signal change due to changing task demands, even if they only differ in the magnitude of their non-oscillatory 1/*f* activity.

When interactions were present in our simulations, dB conversion systematically over- or underestimated the size of the effect as a function of the noise difference between groups, depending on whether the within-subject contrast was larger in the fixed (low broadband activity) or variable noise (increasing broadband activity) group, respectively. In the latter case, the direction of the effect reversed at the most extreme noise-difference levels. This strongly suggests that dB converted results can lead to spurious effects and incorrect conclusions.

Our findings showed that simple baseline subtraction outperformed dB conversion in all scenarios. This is unsurprising given that our data were generated through an additive process. Baseline subtraction was not impervious to the effect of noise, but this noise-dependence only manifested in a decrease in statistical power (*t*-values tending towards 0) as a function of broadband activity difference between groups. This simply reflects the inverse relationship between signal-to-noise ratio and power. No distortions that could have a substantial deleterious effect on the interpretation of findings (e.g., effect reversals, or illusory effects) were observed in the noise range examined.

### Limitations and suggestions for future research

The present study is not without its limitations. Other artifacts, besides the imprecision of 1/*f* slope estimation, could have contaminated our results regarding the relationship between the level of background activity and alpha power. We used a fixed frequency interval based on the canonical alpha frequency band to capture alpha power, but this method does not take into account potential shifts in the central frequency of alpha within participants, across time (Donoghue et al., in press). However, unless these within-subject frequency shifts are correlated with changes in 1/*f* activity – an empirical question in its own right - they are unlikely to systematically distort our findings. There are also multiple methods proposed in the literature of ensuring that the estimation of 1/*f* slope is not contaminated by narrowband oscillations, and vice versa (Donoghue et al., in press; Ouyang et al., 2020; Wen & Liu, 2016). We elected to simply exclude canonical frequency bands from our slope estimation, and subtract the estimated slope from the original spectrum, because this is a computationally efficient method of separating these two sources of activity. Although this method is unlikely to lead to perfect separation in all cases, our simulated findings regarding the correlation between 1/*f* activity and narrowband power suggest that this feature would be unlikely to mask a strong positive correlation that would be predicted by the multiplicative model.

Even in light of these limitations, based on our simulated findings regarding dB conversion, it is clear that the choice of baseline correction method needs to be motivated appropriately in time-frequency analysis studies, and the assumptions behind this decision must be made explicit. If one assumes that the relationship between non-oscillatory broadband activity and oscillatory narrowband activity is additive, a subtractive baseline is the appropriate choice, whereas if the assumption is that a multiplicative relationship exists, baseline division is justified (Grandchamp & Delorme, 2011). Here, we presented some evidence in favor of an additive relationship. Notably, most recent algorithms designed to separate 1/*f* scaling from oscillatory peaks in the frequency domain involve subtracting the former from the amplitude or power spectrum, in line with an assumption of additivity (Donoghue et al., in press; Ouyang et al., 2020; but see Demanuele et al., 2007).

When an additive model was assumed, distortions in the data led to erroneous conclusions regardless of the statistical procedure used to assess significance. Pseudo-confirmatory analyses modelled a situation where researchers have *a priori* hypotheses about the frequency range, timing, and duration of the effect, and were based on mean power in a predefined window, whereas pseudo-exploratory analyses were based on permutation testing with a cluster-based correction for multiple comparisons (Cohen, 2014b), modelling a scenario where researchers are interested in exploring when and where a significant effect occurs. As both of these analyses produced unwanted noise-dependent outcomes, it seems clear that the concerns created by problematic pre-processing cannot be mitigated by the choice of analytic strategy or by having strong *a priori* hypotheses and thereby limiting the focus of analysis to a region of the time-frequency space.

As noted above, these findings are likely to be of high importance for researchers investigating between-group and individual differences, such as cross-sectional studies of development, aging, or psychiatric disorders. Neural variability as captured by characteristics of the 1/*f* slope of the power spectrum is likely to differ between various age groups (Clements et al., 2020; Dave et al., 2018; Voytek et al., 2015) and healthy and clinical populations (e.g., in ADHD; Robertson et al., 2019). In such studies, it might be prudent to ensure that unexpected findings are not the result of pre-processing based on inappropriate assumptions, for example by investigating the correlation between levels of broadband and narrowband activity or by re-processing data using a different baseline correction method and reporting both in the Results section. This could ensure that the generalizability of results across different models can be established. The online scripts associated with this paper contain code for the implementation of both of these steps. One viable option is to pre-register these decisions (e.g., what strategies will be adopted to check the multiplicativity assumption of dB conversion, and which baseline correction strategies will be used depending on the outcome of those checks) prior to analysis to lower the chance of Type I errors (Nosek et al., 2019).

## Conclusion

We presented empirical evidence that the relationship between broadband activity as captured by the offset of 1/*f* scaling in the frequency domain and narrowband alpha power is not consistent with a multiplicative model of signal generation, in which noise and signal scale by the same factor. Then, through a series of simulations, we examined the consequences of using dB conversion, a baseline correction method based on assumptions of multiplicativity, in time-frequency analysis of neural data when signal and noise are in fact additive. A simple mixed-design, common in clinical and life span research, was simulated with a between-subject and a within-subject factor. Our findings showed severe distortions in time-frequency outputs following dB conversion, whereby effects in the high broadband-activity group were systematically attenuated compared to the low broadband-activity group, resulting in the illusory emergence of complex effects even if in reality there were none. As cognitive neuroscientists, we should exercise caution when deciding what kind of baseline correction (subtraction or division based) to use in time-frequency data analysis and make this decision process explicit in reports.

## Acknowledgments

This work was supported by NIA grant RF1AG062666 to G. Gratton and M. Fabiani.

## Footnotes

1 This manipulation alters the offset of the 1/*f* curve in log-log space while leaving its steepness, i.e., the exponent of the curve, unchanged. Shifting the offset means that power at low frequencies and power at high frequencies change in a correlated manner (both increase or decrease), whereas modulating steepness introduces anti-correlation between low and high frequency power (Donoghue et al., in press). This difference is of no consequence to our findings as we did not investigate cross-frequency effects.

2 As the latency of the peak did not vary systematically across conditions or groups in our simulations, this approach is likely to give similar results to an approach where only one fixed time-frequency window is used across all participants. However, in real data, this latter approach might be suboptimal as the latency of an effect could differ across groups or conditions, leading to an inaccurate estimate of conditional power. This is why we decided to set up our simulations using this flexible peak finding method.

3 The eyes open condition showed identical results, see **Supp. Fig. 1**.

4 Grandchamp & Delorme (2011) suggested using unbiased single-trial baseline correction as opposed to baseline correction of the trial-averaged time-frequency matrices, for both subtraction- and division-based methods. This involves first correcting each individual epoch by subtracting from it or dividing it by the mean activity of the whole epoch, not just the baseline period. Then, these shifted epochs are averaged, and the resulting averages are corrected again in the traditional way (subtraction or dB conversion), using mean activity in the baseline period. These methods are also implemented in the online code, but findings are not reported here because they are largely indistinguishable from their trial average-based counterparts.

5 In our simulations, mean baseline activity in a pre-stimulus window was subtracted from all time-frequency points. This is basically the same as estimating 1/*f* and then subtracting that, because in our simulated data there was never anything else in the baseline period other than 1/*f* activity, so the mean baseline activity as a function of frequency was an estimate of 1/*f* activity in and of itself.

