## Supplementary figures and images for "The impact of 1/*f* activity and baseline correction on the results and interpretation of time-frequency analyses of EEG/MEG data: A cautionary tale"

### Supplemental Fig. 1

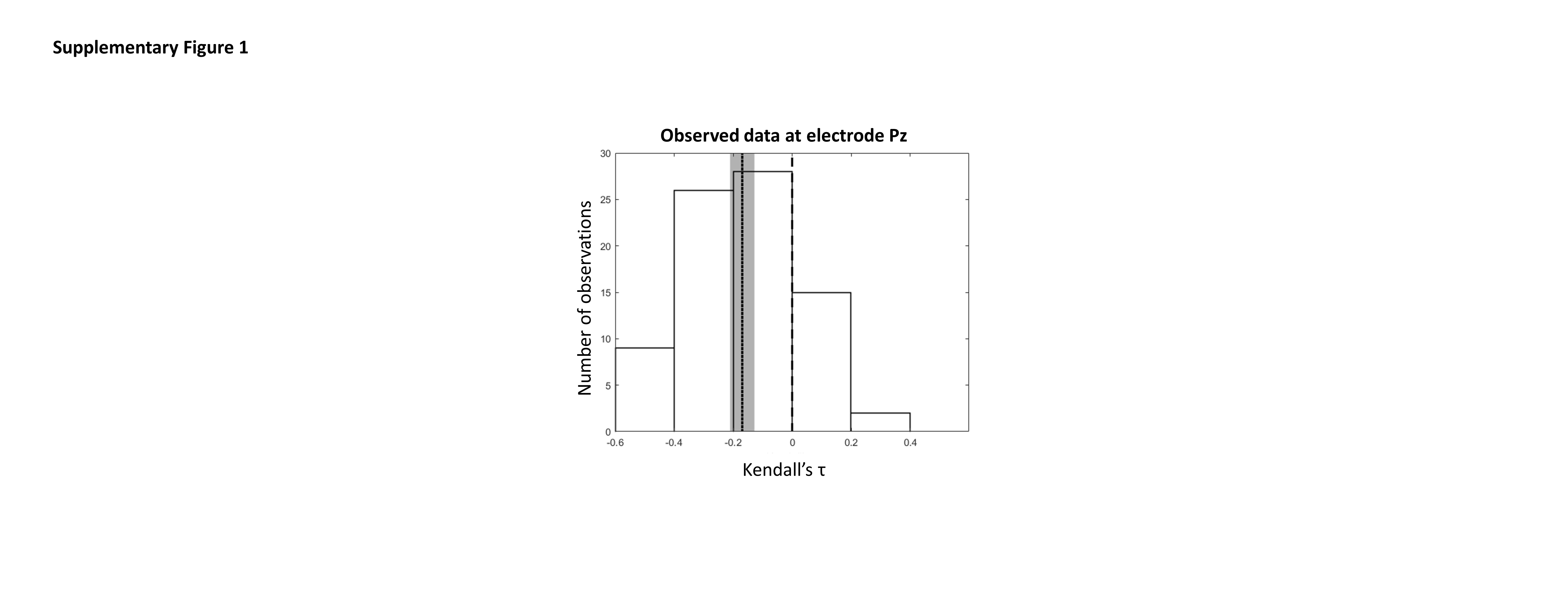
